# Inferring Latent Behavioral Strategy from the Representational Geometry of Prefrontal Cortex Activity

**DOI:** 10.1101/2025.02.19.639166

**Authors:** Yichen Qian, Roger Herikstad, Camilo Libedinsky

## Abstract

Behavioral tasks can be solved employing various strategies. Sometimes, different strategies result in the same observable behavior, making them latent. In this study, we infer the latent behavioral strategy used by monkeys in a working memory updating task by comparing the representational geometry of two prefrontal regions —the lateral prefrontal cortex (LPFC) and the prearcuate cortex (PAC)—with that of recurrent neural network (RNN) models trained to solve the task using different strategies. We found that neural activity patterns in both LPFC and PAC align with only one of the proposed strategies, suggesting that monkeys employ this latent strategy to perform the task. These findings open avenues for investigating the processes that lead to strategy learning and the decision-making mechanisms that determine which strategies are chosen when multiple options are available.

## Introduction

Organisms can respond differently to the same stimulus depending on their internal state, a capability referred to as behavioral flexibility^1^. Additionally, they can use various internal states to produce a similar response to a stimulus, each of which is referred to as a task strategy^2,3^. Different strategies are implemented by different neural dynamics^4–6^ and can be examined mechanistically using recurrent neural network (RNN) models^4,7^. Often, different strategies result in noticeable behavioral differences, such as variations in response accuracy^3^ or reaction times^4^. However, in some instances, different strategies may not produce any observable behavioral differences (i.e., latent strategies).

To date, no studies have been able to infer which latent strategy is used by animals, when multiple strategies are available. Here, we address this gap by comparing the geometric representational properties of two prefrontal regions—the lateral prefrontal cortex (area 9/46, LPFC) and the prearcuate cortex (area 8a, PAC)—with those of RNN models explicitly trained to utilize different strategies to solve the same task.

Monkeys and RNNs were trained to solve a working memory updating task^8^. Human behavioral studies have shown that working memory updating tasks can be performed employing different strategies^9,10^. One possible strategy is to passively store the memory items, and then transfer the information to an active store at the time of recall^11^ (Retrieve at Recall strategy or R@R). Alternatively, another strategy is to rehearse and update the memory online as the items are shown^12^ (Rehearse and Update strategy or R&U). The Retrieve at Recall strategy would presumably involve initially encoding multiple memories in orthogonal neural activity subspaces until one of them is selectively rotated for readout^13^. On the other hand, the Rehearse and Update strategy would presumably involve encoding both the initial and the updated memories directly onto the readout activity subspace.

In both LPFC and PAC we observed activity patterns that were consistent with the RNNs trained to solve the task using the Rehearse and Update strategy, and were inconsistent with the RNNs trained to solve the task using the Retrieve at Recall strategy. Thus, we infer that the monkeys employ the latent Rehearse and Update strategy to solve the task.

## Results

### Behavioral Results

We trained RNNs and two monkeys to perform a working memory updating task (Fig. 1a). In this task, they had to report the location of the last target (red square) seen. The monkeys reported with a saccadic eye movement to the target location. For Target 1/Target 2 (T/T) trials (Fig. 1a top), monkeys had to report the location of Target 2, while for Target 1/Distractor (T/D) trials (Fig. 1a, bottom), monkeys had to report the location of Target 1. The performance of the monkeys, when considering all trials, was higher than 50% (Monkey A: Overall: M(SD) = 0.533(0.011), T/D: M(SD) = 0.506(0.018), T/T: M(SD) = 0.559(0.008); Monkey B: Overall: M(SD) = 0.563(0.036), T/D: M(SD) = 0.563(0.041), T/T: M(SD) = 0.569(0.065)). This is a conservative estimate of performance, since we are including error trails in which the animals made a saccade to the correct target location but were either too slow (> 500 ms), or failed to maintain fixation on the correct target location for longer than 200 ms (Combined: T/D: M(SD) = 0.203(0.109), T/T: M(SD) = 0.158(0.120)). These types of errors, however, do not reflect a lack of understanding of the task, but rather a failure of attention (for responses slower than 500 ms) or a failure of self-control (for when the animals did not maintain fixation on the target for > 200 ms). If we re-categorize these trials as successful, the overall performance would be higher than 60% (Monkey A: T/D: M(SD) = 0.688(0.022), T/T: M(SD) = 0.711(0.032); Monkey B: T/D: M(SD) = 0.626(0.034), T/T: M(SD) = 0.611(0.063)). For subsequent neural data analysis we only included correct trials that were rewarded (i.e., when the animals responded faster than 500 ms and fixated on the correct target for longer than 200 ms). We recorded single neuron activity in areas 9/46 (LPFC) and 8a (PAC) (Fig. 1b). RNNs were trained until they reached >95% accuracy (R@R: M(SD) = 0.998(0.003); R&U: M(SD) = 0.998(0.013)). The inputs to the RNNs could either be Target 1 followed by Target 2 (T/T, Fig. 1c, left), or Target 1 followed by Distractor (T/D, Fig. 1c, right). The R&U strategy was enforced by evaluating the output of the network during both Delays 1 and 2 (Fig. 1d). The R@R strategy was enforced by evaluating the output during Delay 2 only (Fig. 1e).

**Fig. 1.**
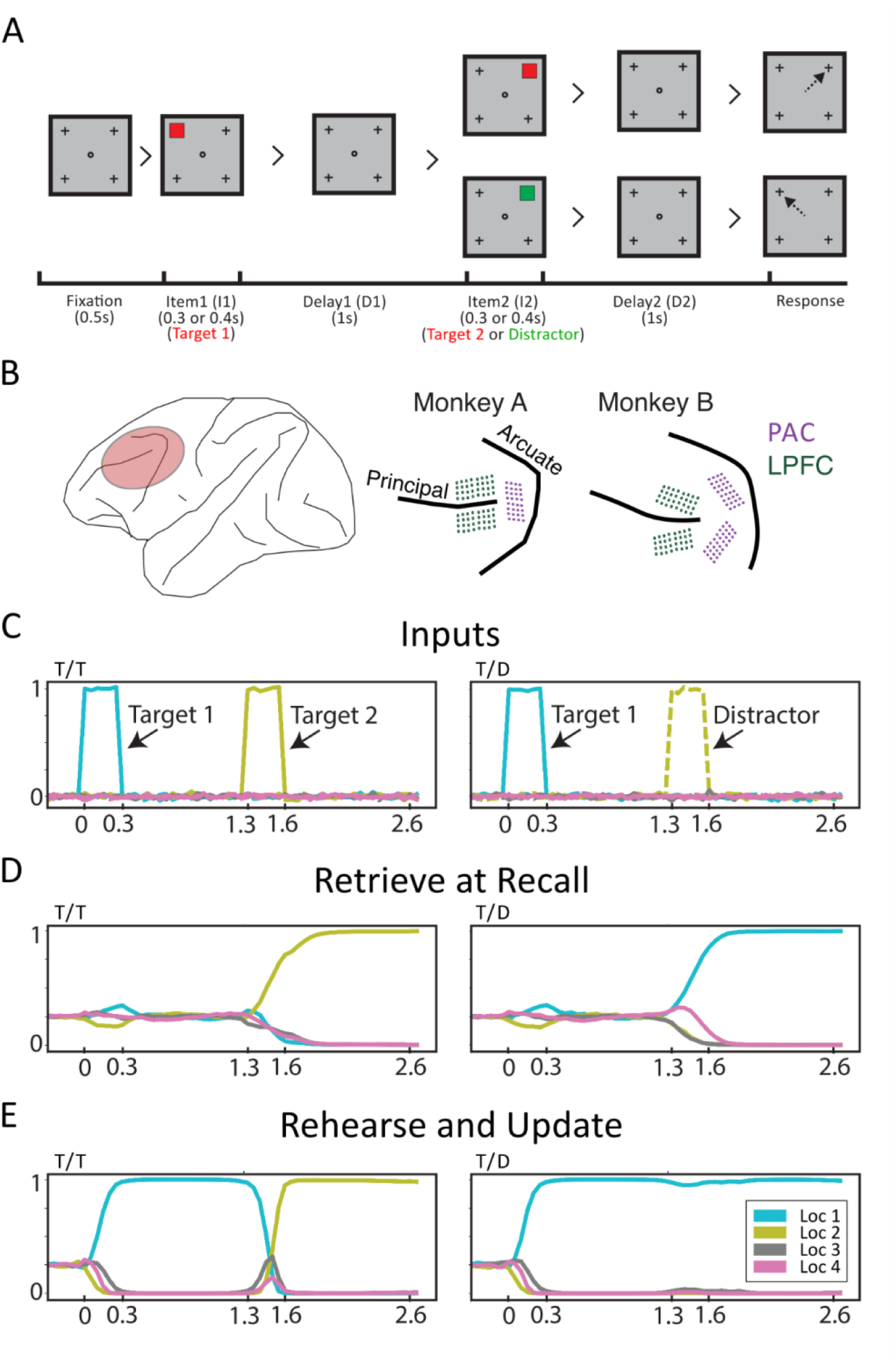
Experiment and model designs. **(A)** Task performed by monkeys. After a 0.5 s fixation period, the Item 1 period always contained a target (Target 1, red square), presented for 0.4 s (Monkey A) or 0.3 s (Monkey B). Following a 1 s Delay 1, an Item 2 period occurred. In T/D trials, Item 2 was a distractor (green square), whereas in T/T trials, it was a new target (Target 2, red square). Item 2 was displayed for 0.4 s (Monkey A) or 0.3 s (Monkey B). After a 1 s Delay 2, the disappearance of the fixation spot served as a go cue. To receive a reward (drop of juice), monkeys had to saccade to the last presented target location (Target 1 in T/D trials, Target 2 in T/T trials). **(B)** Electrode implantation sites. We chronically implanted 64 electrodes in the lateral prefrontal cortex (LPFC) (red-highlighted area) of each monkey, along with either 32 (Monkey A) or 64 (Monkey B) electrodes in the prearcuate cortex (PAC) (purple). **(C)** Input layer activity of RNNs in example T/T (left) and T/D (right) trials. The RNN input layer consisted of 9 units (4 locations × 2 types + 1 fixation). Colors indicate item locations, and line styles indicate item types (solid: target, dashed: distractor). Display durations of Item 1 and Item 2 were fixed at 0.3 s across all RNNs. **(D)** Output activity of Retrieve at Recall (R@R) RNNs in example T/T and T/D trials. The RNN output layer comprised 4 units corresponding to the possible locations. The output units of the R@R RNNs remained uninformative until Item 2 appeared. For T/T trials, the output represented the Item 2 location during Delay 2. For T/D trials, the output corresponded to the Item 1 location. **(E)** Output activity of Rehearse and Update (R&U) RNNs in example T/T and T/D trials. The R&U RNNs exhibited informative output immediately after item presentation. In T/T trials, the output corresponded to the Item 1 location during Delay 1 and switched to the Item 2 location in Delay 2. In T/D trials, the output remained at the Item 1 location across both delays.

### Full Space Decoding Analysis

To get a rough estimate of how information was coded in the models and cortical regions, we carried out a cross-temporal decoding analysis (Fig. 2). First, we evaluated the stability of the code during late Delay 1 and 2 (i.e., the last 500 ms of each delay) (Fig. 2b). To do this we calculated the ratio of the decoding accuracy for time-displaced decoders (i.e., off diagonal) and time-specific decoders (i.e., diagonal). A value of 1 indicates a stable code that does not change across time, while a value lower than 1 reflects a dynamic code. For T/T trials, during Delay 2, both regions and both models showed stable codes (R@R: M(SD) = 0.999(0.003), p = 0.740; R&U: M(SD) = 0.998(0.006), p = 0.640; LPFC: M(SD) = 0.996(0.016), p = 0.730; PAC: M(SD) = 0.995(0.014), p = 0.710) (Fig. 2b, light colors). However, while for the LPFC, PAC and R&U RNNs the code during Delay 1 was equally as stable as during Delay 2, that of the R@R RNNs was more dynamic (R@R: MD = -0.092, p < 0.001, g = -1.863; R&U: MD = -0.001, p = 0.436, g = -0.114; LPFC: MD = -0.001, p = 0.550, g = -0.086; PAC: MD = 0.000, p = 0.766, g = 0.033) (Fig. 2b).

**Fig. 2.**
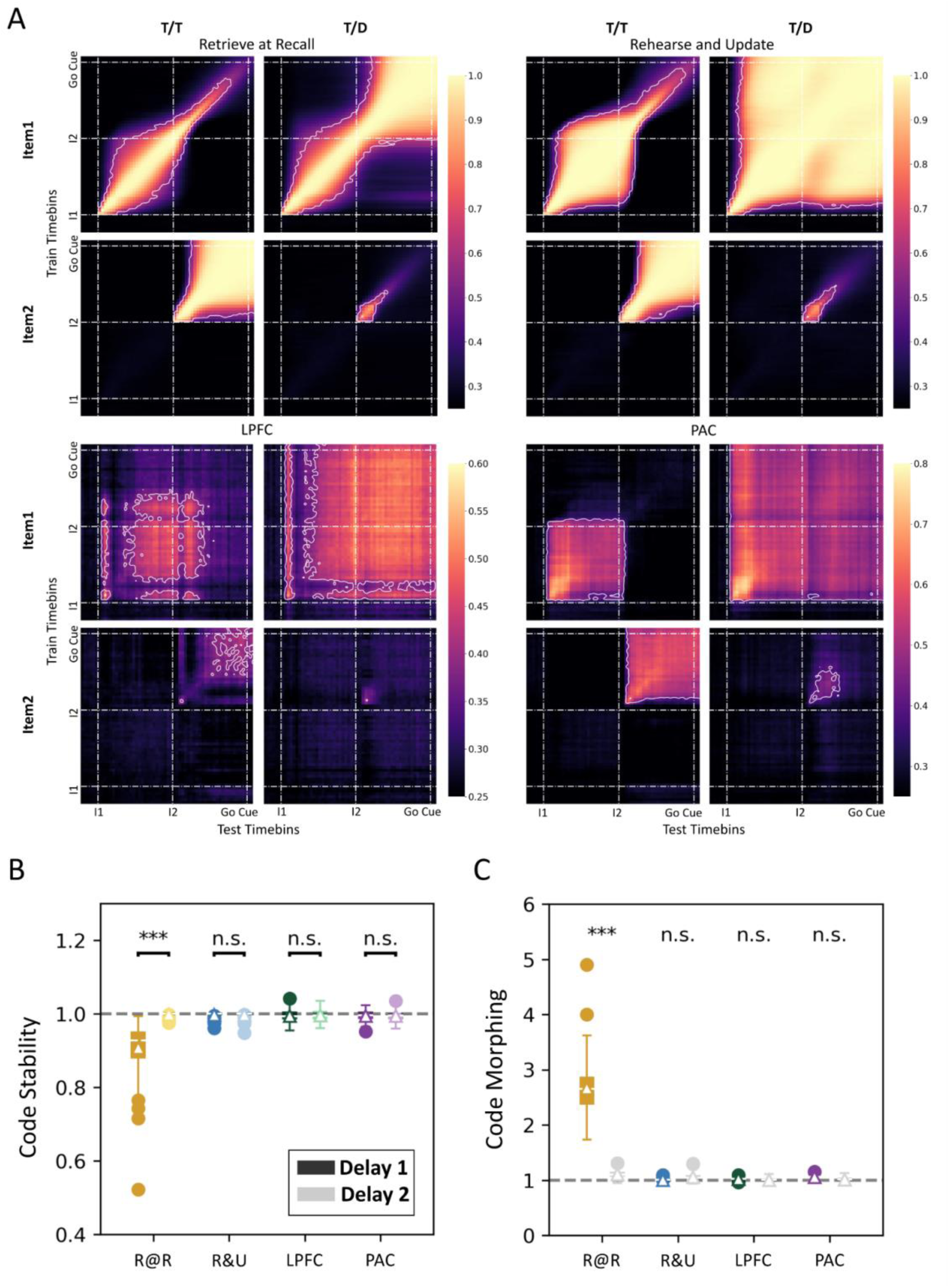
Decodability of item location from the full state space. **(A)** Cross-temporal decodability. Mean decoding accuracy of Item 1 and Item 2 locations from the full state space of R@R (upper left) and R&U (upper right) RNNs, as well as LPFC (lower left) and PAC (lower right) populations. Item 1 (I1), Item 2 (I2), and Go cue onsets are indicated by dotted lines. White outlines indicate significantly above-baseline decodability (p < 0.05). **(B)** Stability ratio distributions. Distributions of target location stability during Delay 1 (dark colors) and Delay 2 (light colors) in R@R (yellow) and R&U (blue) RNNs and LPFC (green) and PAC (purple) populations. **(C)** Code morphing ratio distributions of Item 1 location code across delays in T/D trials, with significance levels compared to corresponding baseline distributions (gray) indicated. In B-C the mean and median values are marked by triangles and white lines, respectively. Significance of the stability ratio difference between Delay 1 and Delay 2 is indicated (***p < 0.001; **p < 0.01; *p < 0.05; n.s.: p > 0.05). Single monkey analyses can be found in Extended Data Fig. 1a-c and Extended Data Fig. 2a-c.

Second, in T/D trials we assessed whether the code for the target location changed between Delays 1 and 2 (i.e., the degree of code morphing^14–16^, Fig. 2c). Target code was generalizable between Delay 1 and 2 in the LPFC, PAC, and R&U RNNs, while it morphed significantly in the R@R RNNs (R@R: M(SD) = 2.670(0.472), p < 0.001; R&U: M(SD) = 1.004(0.011), p = 0.830; LPFC: M(SD) = 1.023(0.023), p = 0.670; PAC: M(SD) = 1.057(0.027), p = 0.420).

### Geometry of Item-Encoding Subspaces

Decoding is a valid, but coarse way of assessing the geometry of the representations in neural networks. To get a more fine-grained assessment of the representational geometry we analyzed the principal angles and alignment of the activity subspaces that encode target information in the two delay periods (see Methods). If two subspaces are identical, then we expect them to have an angle of zero and a perfect alignment, while if they are different (e.g., rotated), then we expect them to have either angles or misalignments larger than zero (Fig. 3a). We also analyzed the transferability of item codes between subspaces (the performance of decoders trained on one subspace and tested on the other), as we expected them to be transferable between identical subspaces, but not so between rotated subspaces.

**Fig. 3.**
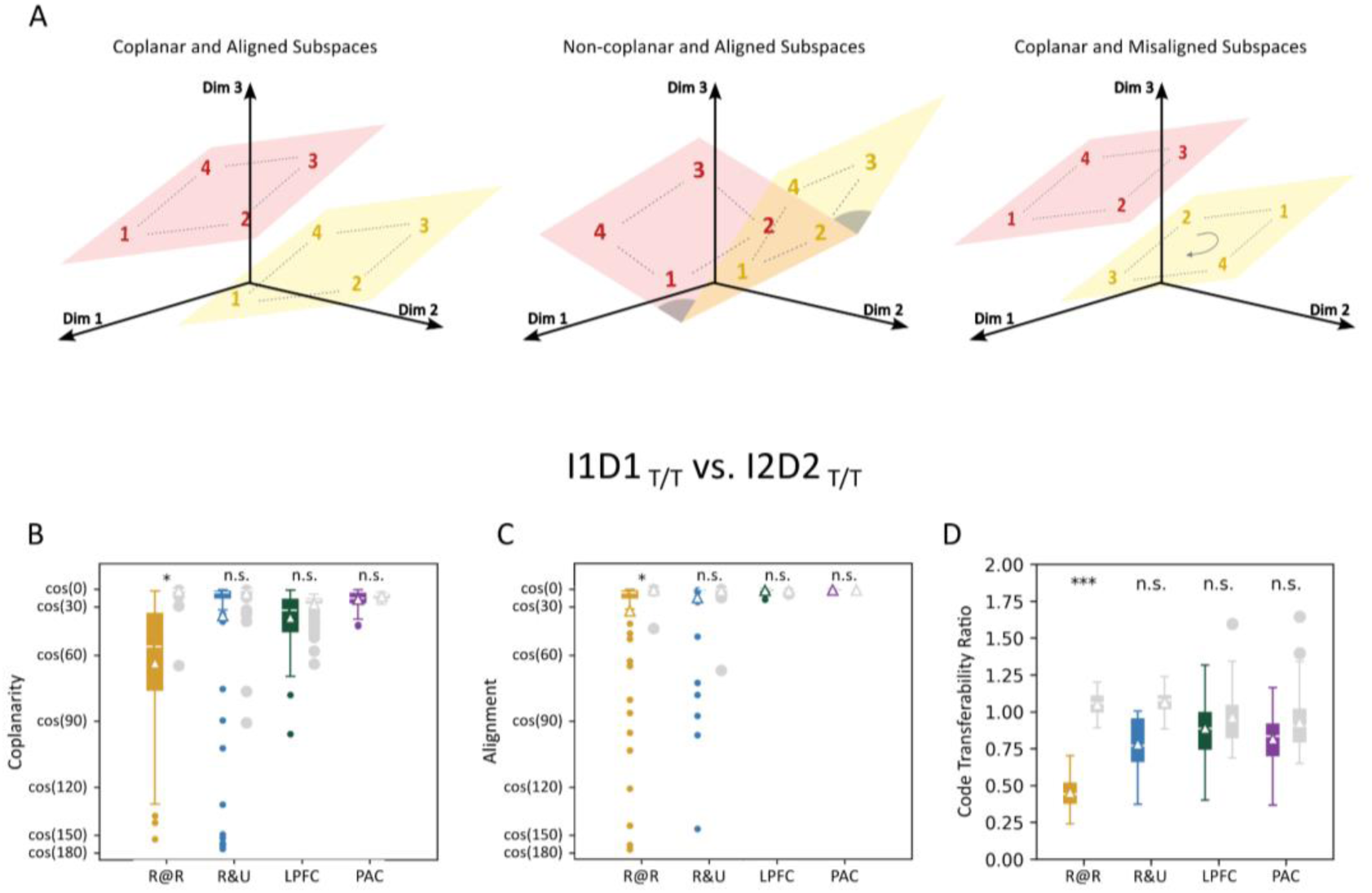
Geometry of item location coding subspaces. **(A)** Schematic of 2D subspaces with varying geometric relationships. Left: subspaces that are coplanar, and with aligned representational configurations. Middle: subspaces that are aligned but non-coplanar. Right: subspaces that are coplanar but misaligned. Subspaces can also be non-coplanar and misaligned (not shown). **(B)** Distributions of coplanarity (cosine of principal angles) between I1D1 T/T and I2D2 T/T subspaces in R@R (yellow) and R&U (blue) RNNs, as well as LPFC (green) and PAC (purple) populations. **(C)** Distributions of subspace alignment (cosine of minimal rotational angles) between I1D1 _T/T_ and I2D2 _T/T_ subspaces, with significance levels compared to corresponding baseline distributions indicated. **(D)** Distributions of code transferability ratios between I1D1 _T/T_ and I2D2 _T/T_ subspaces, with significance levels compared to corresponding baseline distributions indicated. In B-D the mean and median values are marked by triangles and white lines. Significance relative to baseline distributions (gray) is indicated (***p < 0.001; **p < 0.01; *p < 0.05; n.s.: p > 0.05). Single monkey analyses can be found in Extended Data Fig. 1d and Extended Data Fig. 2d.

In T/T trials we estimated the activity subspace that encodes item 1 location during the last 500 ms of Delay 1 (I1D1 _T/T_) and the subspace that encodes item 2 location during the last 500 ms of Delay 2 (I2D2 _T/T_). We found that these subspaces in LPFC, PAC and R&U RNNs were coplanar (cosine of the principal angle was not significantly smaller than 1), while the R@R RNNs they were not coplanar (Fig. 3b) (R@R: M(SD) = 0.446(0.471), p = 0.030; R&U: M(SD) = 0.804(0.493), p = 0.380; LPFC: M(SD) = 0.789(0.194), p = 1.000; PAC: M(SD) = 0.931(0.057), p = 1.000). Similarly, the subspaces in LPFC, PAC and R&U RNNs were aligned (cosine of misalignment angle was not significantly smaller than 1), while the R@R RNNs were not aligned (Fig. 3c) (R@R: M(SD) = 0.839(0.400), p = 0.040; R&U: M(SD) = 0.940(0.253), p = 0.970; LPFC: M(SD) = 0.994(0.010), p = 1.000; PAC: M(SD) = 0.998(0.003), p = 1.000). Finally, we found the code transferability ratio between I1D1 _T/T_ and I2D2 _T/T_ was significantly lower than baseline in R@R RNNs, but not in R&U RNNs, LPFC, and PAC (Fig. 3d) (R@R: M(SD) = 0.453(0.108), p < 0.001; R&U: M(SD) = 0.782(0.172), p = 0.200; LPFC: M(SD) = 0.889(0.179), p = 0.930; PAC: M(SD) = 0.816(0.149), p = 0.750). These results indicate that in T/T trials, the subspace that encodes Target 1 information during during Delay 1 is equivalent to the subspace that encodes Target 2 information during Delay 2 in both brain regions as well as in the R&U model, while in the R@R model they are different.

### Neural Trajectories in the “Readout” Subspace

The analyses above show that there is a common representational subspace in T/T trials during Delay 1 and Delay 2 for the PAC, LPF, and R&U models, but not for the R@R models. To determine whether the same subspace is also utilized during Delay 2 in T/D trials, we compared the same metrics (coplanarity, alignment, and code transferability) between the representational subspaces that encode the reported item information during Delay 2 in T/T and T/D trials. We found that both brain regions and both models had a common representational subspace during Delay 2 in both trial types (*Extended Data* Fig. 3). Thus, a single activity subspace encodes target location information during late Delay 2 in both regions and models, akin to the “readout” subspace reported by Panicello and Buschman (2021)^13^.

In Fig. 2 we used decoding analyses on the full activity space to show that the R&U model and both PAC and LPFC had similar code stability and code morphing properties, while the R@R model behaved differently. Decoding on projections of activity onto the readout subspace led to equivalent conclusions (Extended Data Fig. 4). To get a deeper understanding of the representational properties of the brain regions and models we analyzed the trajectories of the average population activity projected onto the readout subspace (Fig. 4aA). In particular, we focused on the amount of drift observed in the population trajectories between Delay 1 and Delay 2 (Fig. 4b and Extended Data Fig. 5). We found that the drift between delays in the R@R model was similar in T/T and T/D trials (their ratio was close to 1) (Fig. 4b). On the other hand, the drift between delays in the R&U model, PAC and LPFC was primarily present in T/T trials, and almost absent in T/D trials (their ratio was close to 0) (Fig. 4b) (with selected locations: R@R: M(SD) = 0.834(0.154), p = 1.000; R&U: M(SD) = 0.166(0.099), p < 0.001; LPFC: M(SD) = 0.296(0.094), p < 0.001; PAC: M(SD) = 0.183(0.053), p < 0.001). This observation confirms that the R@R model employs different representational subspaces during Delay 1 and Delay 2, consistent with a “passive” storage subspace during Delay 1 that is rotated to a “readout” subspace during Delay 2 (as was observed by Panicello and Buschman, 2021^13^), while the R&U model employs the same subspace in both delays, consistent with the direct loading onto the “readout” subspace during Delay 1. Both prefrontal regions, PAC and LPFC, show a consistent alignment with the R&U model, and consistent differences with the R@R model. Thus, our data strongly supports the conclusion that the monkeys employed the R&U strategy to solve the task.

**Fig. 4.**
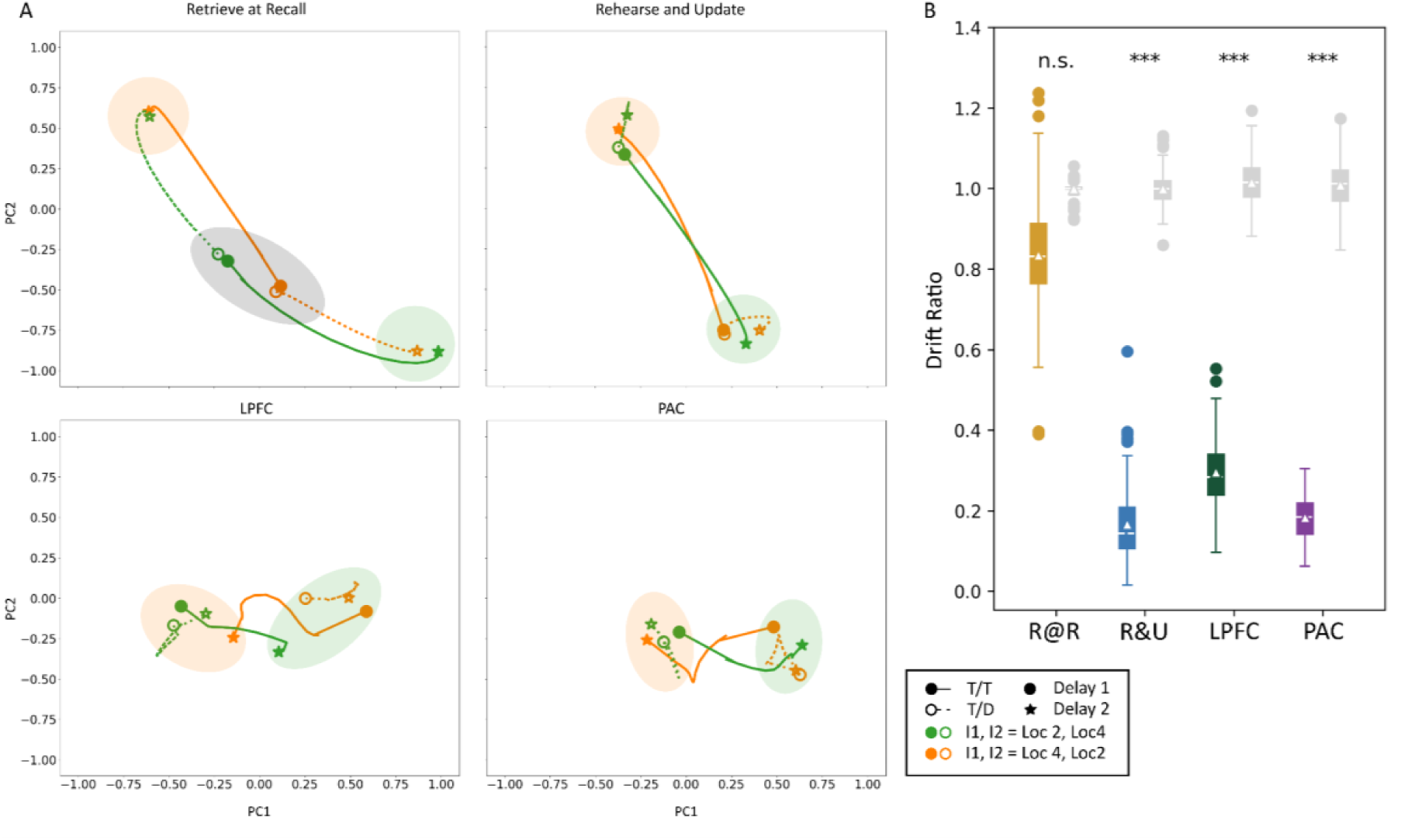
State projections on the readout subspace. **(A)** Population trajectories projected onto the readout subspace. Trajectories of state projections from the end of Delay 1 (circles) to the end of Delay 2 (stars)(green: [I1, I2] = [Loc 2, Loc 4]; orange: [I1, I2] = [Loc 4, Loc 2]) in T/T (solid lines, solid dots) and T/D (dashed lines, hollow dots) trials. Targets in the right hemifield (locations 2 and 4) were selected because they are contralateral to the electrode implantation site - left hemisphere. For the same plots using all 4 locations see Extended Data Fig. 5. Data are shown for example RNNs and neural populations. The approximate representational neighbourhood for the different locations during Delay 2 are highlighted with the large translucent ellipses, for reference only (green: Loc 4; orange: Loc 2). For the R@R network we highlight the region that contains the projections of all target locations during Delay 1 (gray ellipse). **(B)** Distributions of drift ratios for the location pairs in R@R (yellow) and R&U (blue) RNNs, as well as LPFC (green) and PAC (purple) populations. Mean and median values are marked by triangles and white lines. Significance relative to corresponding baseline distributions (gray) is indicated (***p < 0.001; **p < 0.01; *p < 0.05; n.s.: p > 0.05). Single monkey analyses can be found Extended Data Fig. 1e and Extended Data Fig. 2e.

## Discussion

Certain tasks can be solved employing different strategies that lead to *identical* responses to stimuli, in which case the strategy is latent. While previous studies have used behavioral and neural measurements to infer *observable* strategies^4,5,9–12^, to date we have not been able to infer *latent* strategies. Since, by definition, latent strategies cannot be discerned based on behavior, instead we need to rely on differences in neural signals across strategies. In this study, we inferred the latent strategy employed by monkeys by comparing the geometry of their neural representations with the geometry of the representation of RNNs trained to solve the same task employing two different strategies.

Human studies have shown that the running memory span task can be solved using different strategies. In this task, a list of elements is presented in succession for an unpredictable number of items, after which the subjects are asked to recall the last *n* (∼10) items presented^17^. One strategy that can be employed involves storing the items passively, and, when the list ends, attempt to recall as many items as possible from this passive storage^11^. We refer to this strategy as the Retrieve at Recall (R@R) strategy. Panichello and colleagues (2021) used neural data analysis to show that a similar strategy can be used to solve a retrocue task^13^. The authors trained monkeys to retrospectively select one item from a set of items held in short-term memory. They showed that prior to attentional selection, memory items were represented in independent subspaces of neural activity in prefrontal cortex, but after selection these representations were transformed to a “readout” subspace, which was used to guide behavior^13^. Furthermore, Piwek and colleagues (2023) found that this neural transformation emerged naturally in RNNs trained to perform the same task^18^. Our memory updating task can also be solved using this strategy, as shown here using RNNs (Fig. 1d).

Another strategy that can be employed to solve the running memory span task involves keeping track of the last few items, by dropping older ones and adding new ones in an updating process^12^. We refer to this strategy as the Rehearse and Update (R&U) strategy. Our memory updating task can be solved using this strategy, as shown here using RNNs (Fig. 1e). Prefrontal cortex plays an important role in working memory updating^8,19–21^. Here, we showed that monkey prefrontal activity is consistent with the R&U strategy, and inconsistent with the R@R strategy. Thus, using neural signals we were able to infer the latent strategy employed by the monkeys to perform the task. However, this should not be interpreted as a claim that the R&U strategy is the only strategy that monkeys can use, since humans have been shown to employ either strategy under different task conditions^10^. Furthermore, while it has been suggested that monkeys and humans employ different strategies to solve working memory tasks^3^, this discrepancy may be explained by different stages of learning, rather than differences in strategies in both species^7^.

Although this is an important step in understanding how strategies are implemented, our findings do not explain why animals prefer one strategy over another. To address this, we would need to investigate the learning process that leads to the selection of a specific strategy, which may itself involve latent factors^22,23^. Furthermore, when organisms learn that multiple strategies can be used to effectively solve a task, we must explore the decision-making process —both conscious and unconscious^24^— that determines why one strategy is chosen over the others^25^.

## Methods

### Subjects and Surgical Procedures

Two adult male macaques (Macaca fascicularis) were used in this experiment: Monkey A (age 12) and Monkey B (age 12). All animal procedures were approved by, and conducted in compliance with, the standard of the National University of Singapore Institutional Animal Care and Use Committee (NUS IACUC #R18-0295). Procedures also conformed to the recommendations described in Guidelines for the Care and Use of Mammals in Neuroscience and Behavioral Research (National Academies Press, 2003). Each animal was first implanted with a titanium head-post (Crist Instruments, MD, USA) before arrays of intracortical microelectrodes (MicroProbes, MD, USA) were implanted in multiple regions of the left frontal cortex. In Monkey A, two arrays were placed over the dorsal and ventral aspect of the LPFC (Area 9/46), one array was placed over PAC (Area 8A), and one array was placed over the pMA (not included in the analyses here), with 32 electrodes in each. In Monkey B, two arrays of 32 electrodes were placed over the PAC, and two arrays of 32 electrodes each were placed over the LPFC (as shown in Fig. 1b). The arrays consisted of platinum-iridium wires with 200 µm separation, 1-5.5 mm long and with 0.5MΩ of impedance, arranged in 8 x 4 grids. For arrays positioned in the pre-arcuate region (PAC), we used standard electrical microstimulation to confirm that saccades could be elicited with low currents in monkey A (however, we could not perform this procedure in monkey B).

### Recording Techniques

The neural signals in both monkeys were recorded using a 128-channel Grapevine recording system (Ripple Neuro, UT, USA) at 30 kHz sampling rate. The wideband signals were bandpass-filtered between 300 and 3,000 Hz, and spikes were detected on each channel separately using an automated sorting algorithm based on Hidden Markov modelling (Herbst et al., 2008). We recorded the eye positions of each subject using the Eyelink 2000 (SR Research Ltd, ON, CA) on another standalone computer. We designed and ran the behavioral tasks using the Psychopy in Python^26^ on a third computer connected to the recording computer using parallel ports for event mark synchronization during recording.

### Behavioral Tasks

Both monkeys performed a 2-item delayed saccade task. Each trial began with a 0.5 s fixation period, during which the monkey fixated on a white central dot. A first stimulus (Item 1, I1) was then presented at one of four possible locations on the corners of a 3 × 3 grid (10° visual angle) for a fixed duration (Monkey A: 0.4 s; Monkey B: 0.3 s), followed by a 1 s delay (D1) where the monkey maintained fixation. A second stimulus (Item 2, I2) was then presented at one of the three remaining locations for the same duration. I1 was always a task-relevant target (red square), while I2 was either a task-irrelevant distractor (T/D) or a new target (T/T), with equal probability. The target (red) and distractor (green) were identical in shape and size. After a second 1 s delay (D2), the fixation dot disappeared, signaling the monkey to make a saccadic response to the location of the most recent target (I1 in T/D trials; I2 in T/T trials). In other words, the animal needed to make an eye movement to the location of I1 when I2 was a distractor, or to the location of I2 when it was a new target. A juice reward was delivered for correct responses within 0.5 s after the go cue. I1 and I2 never appeared at the same location within a trial, yielding 24 trial conditions based on I1 × I2 locations (4 × 3) and I2 types (2).

### Neuron Firing Rates and Preprocessing

Single-neuron firing rates were computed using a 50 ms sliding window (10 ms overlap). Firing rates were z-scored relative to the mean and standard deviation of pre-I1 baseline activity (0.2 s) across all trials^15^. Since I1 and I2 display durations differed between Monkey A (0.4 s) and Monkey B (0.3 s), we removed the middle 0.1 s of each delay period in Monkey A for pooled analyses to ensure temporal alignment.

### Pseudo Population

To measure the decodable stimulus location information across the population of recorded neurons, we used linear discriminant analysis (LDA) based on the algorithm from scikit-learn^27^ to predict the locations of Items 1 and 2 (Fig. 2 and Extended Data Fig. 4). We created pseudo populations by sampling equal numbers of trials per condition from each session of both monkeys and pooling these trials together, as if these neurons responded together in each pseudo trial. We created 100 pseudo populations separately for the LPFC (89 neurons) and the PAC (129 neurons). Only correct trials were included in the pseudo population. All neural data analyses were conducted on the pseudo populations.

### Models

We trained artificial recurrent neural networks (RNNs) to simulate neural activity under different latent strategies. Each RNN comprised 9 input units (4 locations × 2 types + 1 fixation), 64 recurrent units, and 4 output units (4 locations). The recurrent dynamics followed:

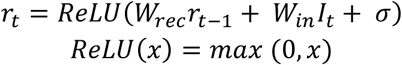

And the output dynamics can be described as:

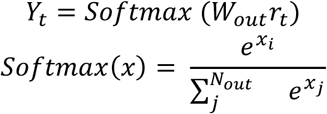

where ***I*** _t_, ***r*** _t_ and ***Y*** _t_ denote the input, recurrent, and output unit activities at time *t*, and ***W*** _in_, ***W*** _rec_, ***W*** _out_ represent the input, recurrent and output weights respectively. ***σ*** represents the noise at the recurrent layer and was drawn from N(0, 0.1). Recurrent weights were orthogonally initialized, while input and output weights followed Xavier-normal initialization. We optimized ***W*** _in_, ***W*** _rec_, ***W*** _out_ using stochastic gradient descent with NAdam. The loss was computed as the mean squared error (MSE) of the outputs over the target time window:

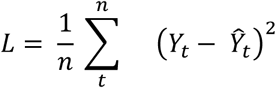

For Retrieve at Recall (R@R) RNNs, during Delay 1 all output units were expected to have equal activity (0.25), and during Delay 2 the output of the choice item (I1 in T/D trials, I2 in T/T trials) was expected to be 1, while the non-choice items remained at 0. For Rehearse and Update (R&U) RNNs, the output corresponded to I1’s location during Delay 1 and switched to the choice item’s location during Delay 2. 100 RNNs were trained per strategy, each with different weight initializations.

### Principal Component Analysis (PCA)

We applied principal component analysis (PCA) to each pseudo-population to reduce dimensionality before decoding, using the scikit-learn library^27^. To preserve variance across conditions, we computed mean firing rates per condition, averaging over Delay 1 and Delay 2. The resulting activity matrix ***X*** (N_conditions_ (24) × N _neurons_) was used to fit a PCA model, which was then applied to the full time series for dimensionality reduction. We defined the full space of the transformed data as the projections on the first 15 PCs, which counted for at least 90% of the total explained variance (EVR) on average in both regions (ΣEVR _PC1-15_: LPFC: M(SD) = 0.908(0.005), PAC: M(SD) = 0.920(0.004)). The same procedure was applied to the hidden unit activities of task-optimized RNNs (ΣEVR _PC1-15_: R@R: M(SD) = 1.000(0.000), R&U: M(SD) = 1.000(0.000)).

### Decoding Analysis

#### Cross-temporal Decodability

Decoding analyses were conducted on neural data from 0.2 s before Item 1 onset to the go cue. A 50 ms non-overlapping smoothing window was applied before training and testing decoders. PCA-based dimensionality reduction was applied before decoding, and analyses were performed on projected activities. The same approach was used for RNN hidden unit activities. For full-space decodability (Fig. 2), we used projections on the top 15 PCs to train and test decoders. For readout subspace decodability (Extended Data Fig. 4), we used state projections on the readout subspace.

Within each pseudo population and RNN, we randomly sampled half of the trials for training and the other half for testing, iterating 100 times. At each iteration, a decoder was trained using training set activities at one time bin to predict item locations and tested on the test set activities across all time bins. Decoding performance was averaged over 100 iterations to minimize variability from set-splitting. We tested decodability significance at each time bin using a 95th-percentile test (see *Statistical Tests*), comparing decoder accuracy to baseline distributions generated from data with shuffled location labels (shuffled 100 times per pseudo-population and RNN). We repeated this process until all time bins were trained and tested.

#### Code Stability & Code Morphing

To quantify information coding stability, we computed the stability ratio, defined as the mean within-time decodability (trained and tested at the same time bin) divided by the mean cross-time decodability (trained and tested at different time bin) across selected time windows^16^. We calculated this ratio separately for late Delay 1 (last 0.5 s) and late Delay 2, followed by statistical comparisons (see *Statistical Tests*).

The population code of the target may change after the presentation of a distracting stimulus (i.e., code morphing^14^). Here we quantified the code morphing ratio in the T/D trials as the mean cross-temporal decodability trained and tested in late Delay 1, divided by the decodability trained in late Delay 1 and tested in late Delay 2 (and vice versa). A low code morphing ratio indicates a time-invariant code. Baseline distributions were created by decoders trained with label-shuffled data, iterated 100 times and averaged within each pseudo-population and RNN.

### Subspace Analysis

#### Subspace Estimation

Low-dimensional geometries can be estimated from the population based on various methods^13,15,18,28^. We estimated low-dimensional subspaces using methods adapted from Panichello & Buschman (2021)^13^ and Piwek et al. (2023)^18^ to obtain 2D planes best fitting the representations of Item 1 and Item 2 during early (first 0.5s of) and late (last 0.5s of) Delay 1 and Delay 2. For each pseudo population, we first grouped the trials by conditions (N _conditions_ = N _location combinations_ (12) × N _type_ (2)). We then averaged the neural population states by conditions and projected the N _conditions_ (24) × N _neurons_ data to the axes of the top three principal components (PCs). On average, the sum of explained variances by these three PCs reached about 50% of total variances in LPFC and over 60% in PAC as well as in R@R and R&U RNNs (ΣEVR _PC1-3_: R@R: M(SD) = 0.995(0.010), R&U: M(SD) = 0.994(0.007); LPFC: M(SD) = 0.499(0.016), PAC: M(SD) = 0.611(0.013)).

Within each time window, we first averaged the activities across all timepoints, returning us an N_conditions_ (24) × N_PCs_ (3) matrix. For item-specific subspaces (Fig. 3; Extended Data Fig. 3), we separated the matrix by the type condition (T/T & T/D) for analysis clarity, each with a 12 × 3 matrix. Within each type, we estimated the mean representation by averaging the rows corresponding to conditions with the same location of each item: for example, the representation of I1 presented at location 1 was calculated as the mean of all conditions with I1-location = 1, regardless of the location of I2. This returned us an N _locations_ (4) × N_PCs_ (3) matrix for each item. Next, we conducted a secondary PCA on this matrix and extracted the first two PCs. These two eigenvectors corresponded to the directions of maximum variances by locations, hence were used to define the best-fitting 2D plane of the locational representation. Finally, we projected the states from the 3-PC space onto these 2D planes and used these projections for subsequent analyses.

The “readout” subspaces (Fig. 4) were estimated based on the same approach, except that we did not separate between trial types, and the mean representation was calculated by averaging the rows with the same location of the final choice item (i.e., I1 for T/D and I2 for T/T). Also, instead of estimating at separate time windows, we only used activities during late D2 to estimate the “readout” axes, as stable outputs were observed during this period in both models and brain regions (as shown in Fig. 2).

#### Geometry Analysis

With the subspace estimated using the above-described approach, we then examined the geometric relationships between subspace pairs, specifically the coplanarity and the degree of alignment of the representational configurations between subspaces. Metrics were calculated using methods adapted from Panichello and Buschman (2021)^13^ and Piwek et al. (2023)^18^. For coplanarity, we measured the cosine of the principal angle between the normal vectors of the two planes. We applied a correction to the directions of the plane-defining vectors based on the representational configurations to guarantee vectors from two different subspaces were under a common frame of reference, as described below^18^. This correction was necessary since the principal angle was sensitive to the normal vector directions, which were essentially determined by the directions of the defining vectors of the 2D plane; yet each eigenvector estimated from PCA (used as the plane-defining vectors) could be oriented to either direction along its PC axis, thus the directions of eigenvectors estimated from separate PCAs might be inconsistent and could lead to mismatched normal vector directions and consequently distorted principal angles. To avoid this problem, we corrected the directions of the eigenvectors by first rotating PC1 vector to be positively collinear (i.e. parallel and in the same direction) to a predefined directional side of the location configurations, creating a PC1_corrected_, and then rotating PC2 vector by the shortest angular distance to be orthogonal to PC1_corrected_. We then used PC1_corrected_ and PC2_corrected_ as the plane-defining vectors as well as for normal vector calculations. Planes with mirror-reflected configurations of one another would lead to angles greater than 90° (cos(θ) < 0), whereas the planes of rigid rotation of each other would have principal angles restricted smaller than 90° (cos(θ) > 0).

For alignment, we measured the cosine of the minimal rotation angle minimizing the distances between corresponding representational points on the two subspaces. To calculate the rotation angle, we first forced the two planes to be coplanar. Next we estimated the representational configuration from each plane by averaging the states by item locations and then projected the configuration points on corresponding planes, giving us two N _locations_ (4) × 2 configuration matrices. We then applied an orthogonal Procrustes analysis on these two matrices, which returned an orthogonal matrix ***R*** that could rotate one matrix to most closely match the other, with the ***R****(0,0)* as the cosine of the minimal rotation angle.

For both coplanarity and alignment, the baseline hypothesis was the full equivalence between subspaces, in other words, we tested whether the two subspaces were significantly non-coplanar and misaligned. To create the baseline distributions, we first split the data into a training set and a test set and separately estimated subspaces. We then calculated the coplanarity and alignment between the training and test set subspaces at each item and time bin of interest and finally took the mean of them. The rationale was that a given subspace is perfectly coplanar and align to itself, and subspaces estimated from different subsets of the same pseudo-session or recurrent neural network (RNN) should be highly similar, with some variability due to differences between training and test sets. For example, for the comparison between I1D1 and I2D2, we averaged coplanarity and alignment between I1D1 _train_ vs. I1D1 _test_ and I2D2 _train_ vs. I2D2 _test_. For each pseudo population and RNN we repeated this process for 100 times and took their average. The resulting parallel distributions were then compared to observed values, and empirical p-values were calculated using the 95^th^-percentile test (see *Statistical Tests*), and a significant p value indicated consistent deviation from the equivalent hypothesis.

#### Code Transferability Ratio

In addition to the geometry analysis, we also examined the code transferability between subspaces. Specifically, we trained LDA decoders with the state projections on one subspace and tested with the projections on the other subspace. The codes were expected to be transferable if the two subspaces were equivalent. Given the noisy nature of neural data, the decoder performances in the LPFC and PAC were generally worse than in the RNNs. To make the quantifications comparable between neural data and RNNs, we calculated the ratio of cross-subspace decodability (i.e., code transferability) over the within-subspace decodability (i.e., trained and tested with the projection on the same subspace), and a high ratio (close to 1) indicates the decodability to be at a similar level cross- and within-subspace. The baseline distributions were created by training the cross- and within-subspace decoders on data with shuffled location labels, and for each pseudo population and RNN we repeated the process for 100 times and took the average.

#### Drift Ratio

To quantify the dynamics of population activity projection onto the readout subspace, for each condition (I1-I2 Combination × Type) we calculated the Euclidean distance between the mean projection points during late (last 0.5 s of) Delay 1 and during late Delay 2 and scaled to the range from -1 to 1. We then calculated the drift ratio by dividing the drifted distance in T/D trials by the T/T trials. A low ratio would indicate the readout projections to be changed only when information updates were required. The baseline distributions were created by estimating the distances from data with shuffled location labels, and we repeated this process for 100 times and took the average for each pseudo population and RNN.

### Significance Tests

For tests comparing to baselines, we considered two distributions to be significantly different from baseline if the 95th percentile range of the test distribution did not overlap with the baseline distribution^29^. The p value was calculated as:

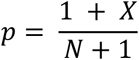

where *X* represents the number of overlapping points and *N* represents the total number of points in the baseline distribution.

For tests comparing between two test distributions, we pooled the data together and randomly reassign them to create two new distributions and calculated the mean difference. We iterated this process for 1000 times to create a null distribution. The p value was then calculated as the proportion of points in the null distribution that are larger or smaller (depending on the test direction of interest) than the true mean difference between the test distributions. We additionally calculated the effect size of the comparisons using the measure of Hedge’s g as:

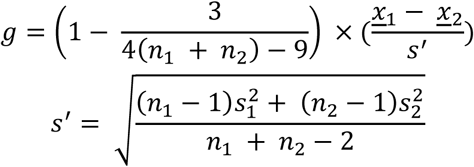

where n and s refer to the number of data points and standard deviations of each distribution^30^.

## Supplementary Material

**Extended Data Fig. 1.**
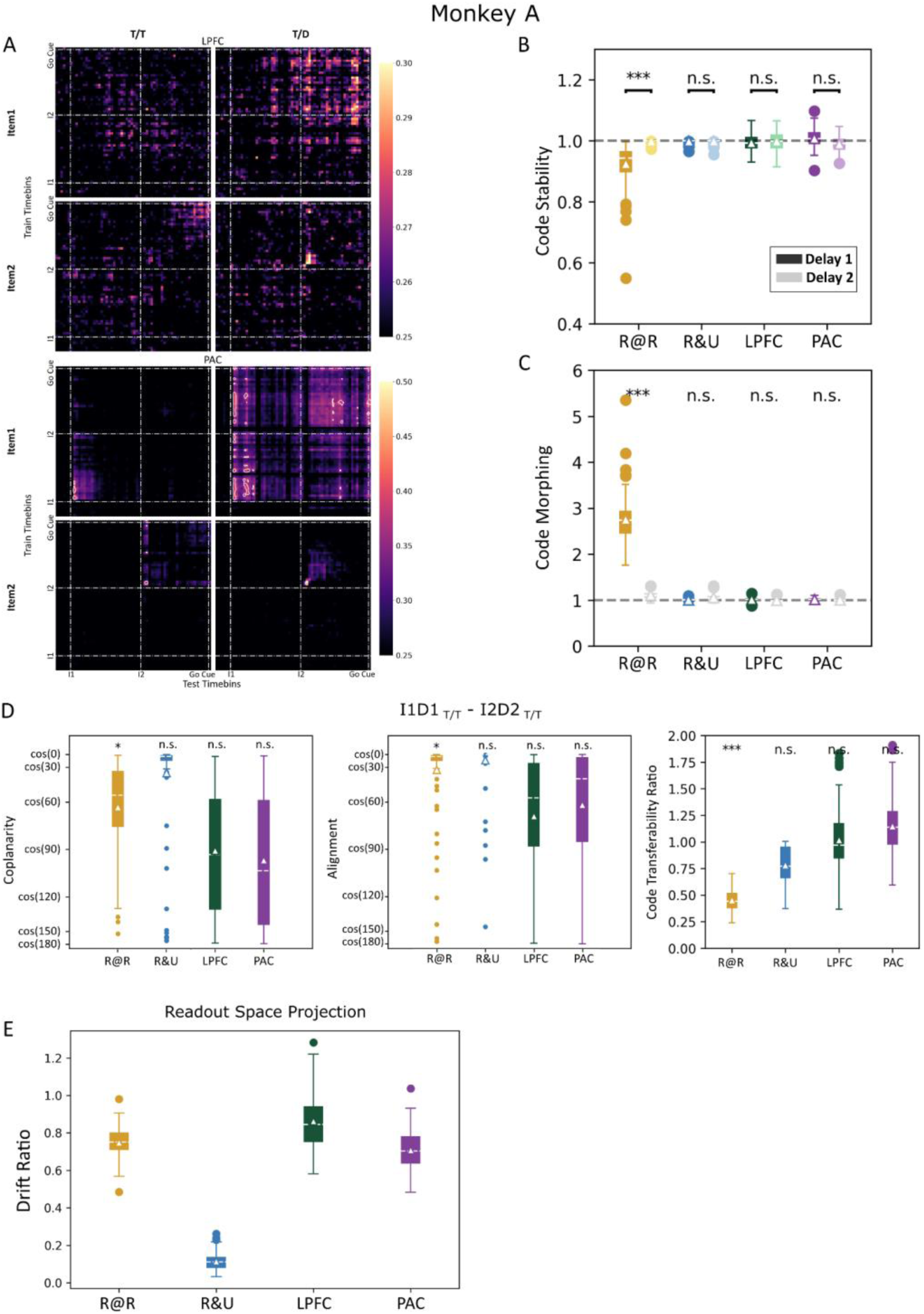
Results for monkey A. **(A-C)** Replication of results shown in Fig. 2a-c. **(D)** Replication of results shown in Fig. 3b-d. **(E)** Replication of results shown in Fig. 4b.

**Extended Data Fig. 2.**
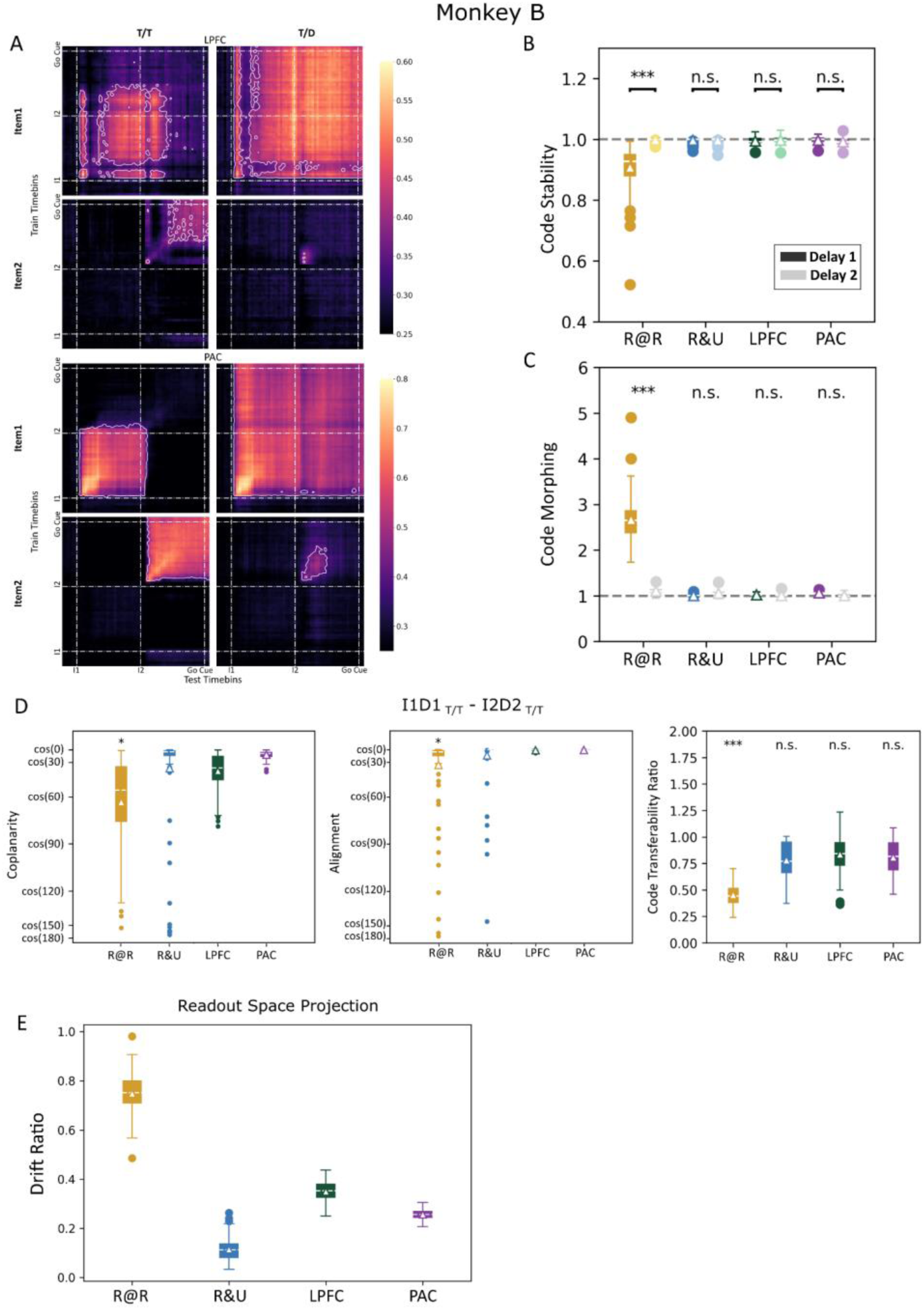
Results for monkey B. **(A-C)** Replication of results shown in Fig. 2a-c. **(D)** Replication of results shown in Fig. 3b-d. **(E)** Replication of results shown in Fig. 4b.

**Extended Data Fig. 3.**
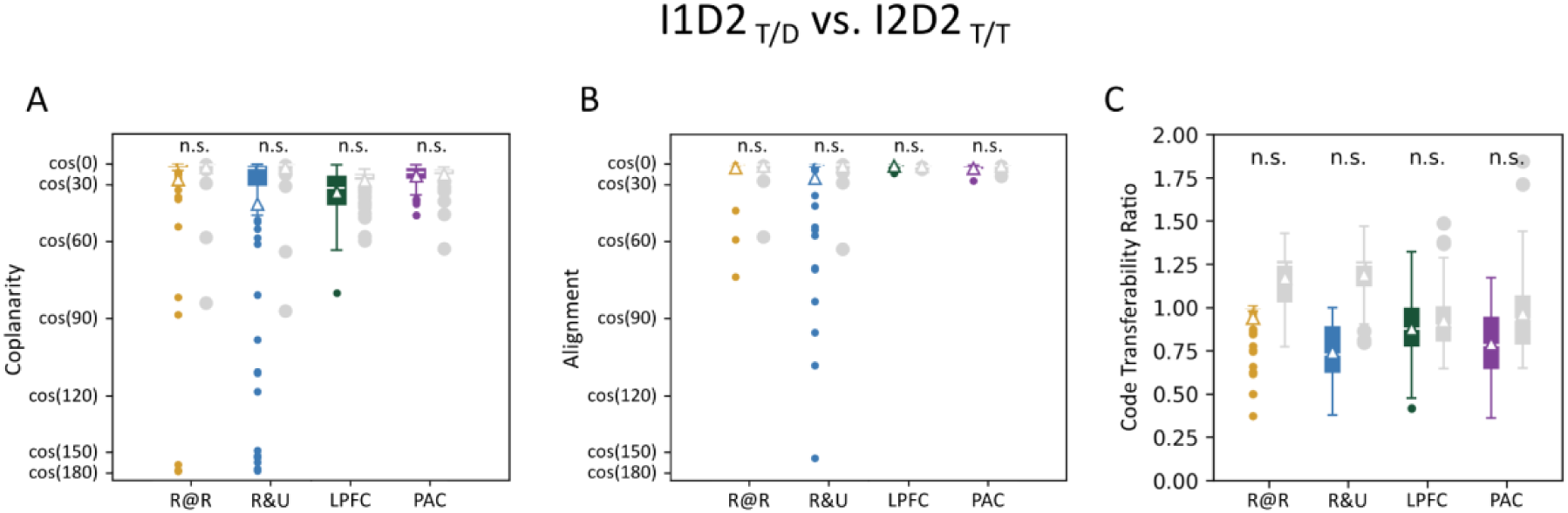
Readout subspace is equivalent in T/D and T/T trials. **(A)** Distributions of coplanarity (cosine of principal angles) between I1D2 _T/D_ and I2D2 _T/T_ subspaces in R@R (yellow) and R&U (blue) RNNs, as well as LPFC (green) and PAC (purple) populations. Mean and median values are marked by triangles and white lines. (R@R: M(SD) = 0.903 (0.357), p = 0.700; R&U: M(SD) = 0.744 (0.534), p = 0.390; LPFC: M(SD) = 0.819 (0.147), p = 1.000; PAC: M(SD) = 0.928 (0.074), p = 1.000). **(B)** Distributions of subspace alignment (cosine of minimal rotational angles) between I1D2 _T/D_ and I2D2 _T/T_ subspaces, with significance levels compared to corresponding baseline distributions indicated. (R@R: M(SD) = 0.984 (0.091), p = 0.960; R&U: M(SD) = 0.915 (0.285), p = 1.000; LPFC: M(SD) = 0.993 (0.010), p = 1.000; PAC: M(SD) = 0.980 (0.013), p = 0.540). **(C)** Distributions of code transferability ratios between I1D2 _T/D_ and I2D2 _T/T_ subspaces, with significance levels compared to corresponding baseline distributions indicated. (R@R: M(SD) = 0.941(0.132), p = 0.230; R&U: M(SD) = 0.741(0.173), p = 0.150; LPFC: M(SD) = 0.877(0.183), p = 0.900; PAC: M(SD) = 0.791(0.186), p = 0.800). For all plots, the significance relative to baseline distributions (gray) is indicated (***p < 0.001; **p < 0.01; *p < 0.05; n.s.: p > 0.05).

**Extended Data Fig. 4.**
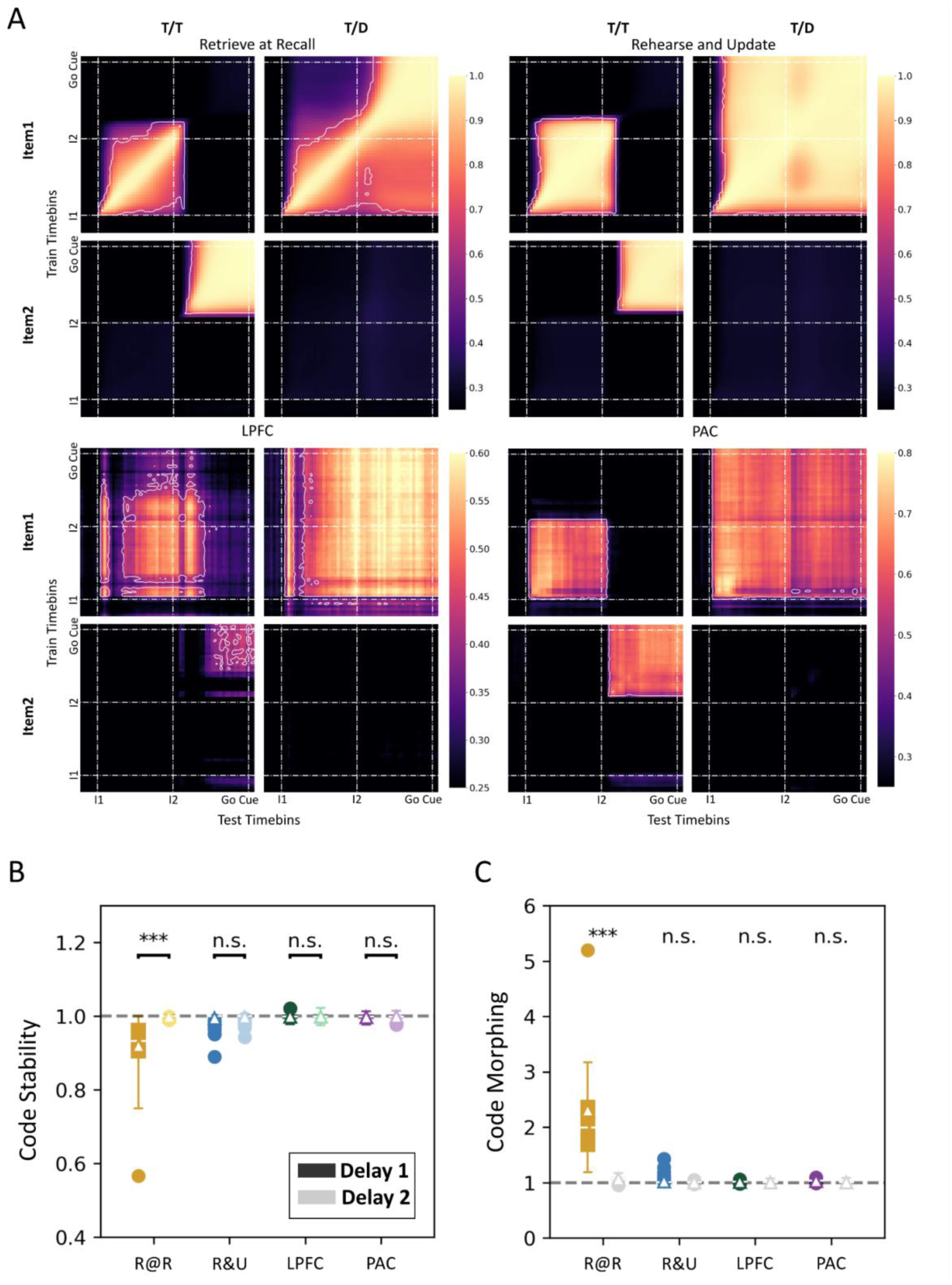
Cross-temporal decoding analysis on readout subspace projections. **(A)** Cross-temporal decodability. Mean decoding accuracy of Item 1 and Item 2 locations from the state projections on the readout subspace of R@R (upper left) and R&U (upper right) RNNs, as well as LPFC (lower left) and PAC (lower right) populations. Item 1 (I1), Item 2 (I2), and Go cue onsets are indicated by dotted lines. White outlines indicate significantly above-baseline decodability (p < 0.05). **(B)** Stability ratio distributions. Distributions of target location stability during Delay 1 (dark colors) and Delay 2 (light colors) in R@R (yellow) and R&U (blue) RNNs and LPFC (green) and PAC (purple) populations. (R@R: MD = -0.079, p < 0.001, g = -1.497; R&U: MD = -0.003, p = 0.086, g = -0.243; LPFC: MD = -0.001, p = 0.702, g = -0.055; PAC: MD = -0.000, p = 0.962, g = -0.012) **(C)** Morphing ratio distributions of Item 1 location code across delays in T/D trials, with significance compared to corresponding baseline distributions (gray) indicated. (R@R: M(SD) = 2.302(2.379), p < 0.001; R&U: M(SD) = 1.019(0.057), p = 0.680; LPFC: M(SD) = 1.012(0.015), p = 0.650; PAC: M(SD) = 1.034(0.017), p = 0.450). For B-C the mean and median values are marked by triangles and white lines, respectively, and the significance of the stability ratio difference between Delay 1 and Delay 2 is indicated (***p < 0.001; **p < 0.01; *p < 0.05; n.s.: p > 0.05).

**Extended Data Fig. 5.**
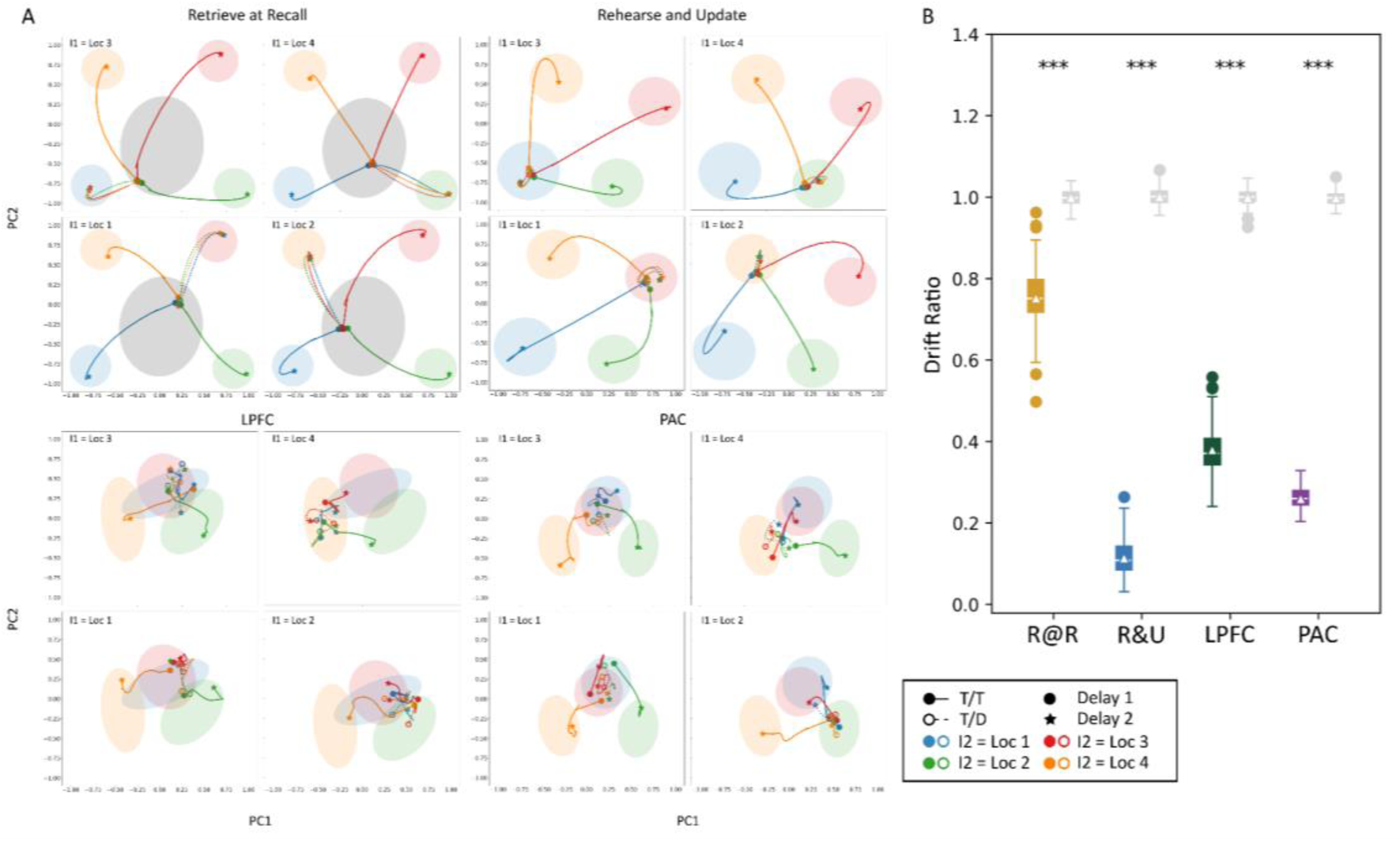
State projections on the readout subspace with all location pairs. **(A)** Population trajectories projected onto the readout subspace. Trajectories of state projections from the end of Delay 1 (dots) to the end of Delay 2 (stars) in T/T (solid lines, solid dots) and T/D (dashed lines, hollow dots) trials with all I1-I2 location pairs. Data are shown for example RNNs and neural populations. I1 locations were labelled on the upper left corner of each panel. I2 locations were color-denoted (blue: Loc 1; green: Loc 2; red: Loc 3; orange: Loc 4), and approximate representational loci of locations are circled in corresponding colors. For R@R networks, approximate range of intermediate saddle points is also highlighted (gray). **(B)** Distributions of drift ratios with all location pairs in R@R (yellow) and R&U (blue) RNNs, as well as LPFC (green) and PAC (purple) populations. (R@R: M(SD) = 0.752(0.078), p < 0.001; R&U: M(SD) = 0.114(0.048), p < 0.001; LPFC: M(SD) = 0.380(0.059), p < 0.001; PAC: M(SD) = 0.259(0.030), p < 0.001). Mean and median values are marked by triangles and white lines. Significance relative to corresponding baseline distributions (gray) is indicated (***p < 0.001; **p < 0.01; *p < 0.05; n.s.: p > 0.05).

